# Epigenetic reprogramming ameliorates type 1 diabetes by decreasing the generation of Th1 and Th17 subsets and restoring self-tolerance in CD4^+^ T cells

**DOI:** 10.1101/2021.07.24.453657

**Authors:** Vasu Patel, Arathi Jayaraman, Sundararajan Jayaraman

## Abstract

The histone modifier Trichostatin A (TSA) ameliorated diabetes and repressed IFN-γ and IL-17A expression in prediabetic female NOD mice. Purified CD4^+^ cells could be polarized *ex vivo* into Th1 and Th17 subsets, which comparably transferred diabetes into NOD.*scid* mice. Polarized Th1 cells were devoid of IL-17A-producing cells and did not transdifferentiate into Th17 cells in an immunodeficient environment. However, Th17 cells had contaminant Th1 cells, which expressed IFN-γ upon adoptive transfer into lymphopenic recipients. Notably, TSA treatment abrogated the transfer of diabetes by CD4^+^ T-cells cultured under Th1 or Th17 polarizing conditions accompanied by the absence of *Ifng* and *Il17a* expression in NOD.*scid* recipients. Significantly, the histone modifier restored the ability of CD4^+^ but not CD8^+^ T-cells to undergo CD3-mediated apoptosis *ex vivo* in a caspase-dependent manner. Thus, the histone modifier afforded protection against autoimmune diabetes by negative regulation of signature lymphokines and restitution of self-tolerance in CD4^+^ T cells.

## INTRODUCTION

Type 1 diabetes (T1D) occurs at an alarming rate in recent years (Gale, 2002). Since concordance among monozygotic twins is less than 50%, environmental factors including microbial infections, diet, and intestinal microbiota have been attributed to variations in diabetes incidence. Female non-obese diabetic (NOD) mice are considered a reliable model for human autoimmune diabetes (Anderson and Bluestone, 2005; Mullen, 2017). Since T-cells are critical for T1D, the nature of the T-cells involved, their specificity, activation requirements, stability, and the mechanisms by which they mediate the disease are under intense investigation. A previous study utilizing the T cell receptor (TCR) transgenic BDC2.5 mice indicated that the CD4^+^ T cells polarized to become interferon-γ (IFN-γ)-producing Th1 cells could adoptively transfer diabetes in neonatal NOD mice (Katz et al., 1995). However, the IL-17A-expressing Th17 cells derived from the BDC2.5 mice could transfer diabetes into NOD.*scid* mice only *after* conversion into Th1 cells in the immunodeficient NOD.*scid* mice (Martin-Orozco et al., 2009; Bending et al., 2009). On the other hand, the use of the IL-17 reporter mice in an experimental autoimmune encephalomyelitis (EAE) model suggested that the Th1 and Th17 cells are *not* terminally differentiated and equally pliable (Kurschus et al., 2010). These studies are also at variance with the view that the Th17 cells can develop via a lineage distinct from the Th1 and Th2 type cells (Harrington et al., 2005). These data raise the question of whether the CD4^+^ T cells from unmanipulated *wild-type* NOD mice could also be similarly polarized to become distinct Th1 and Th17 subsets and mediate diabetes independently of each other.

Clonal deletion of immature autoreactive thymocytes is a crucial tolerance mechanism for ensuring self-tolerance, and its impairment can lead to autoimmune diseases (Ohashi, 2003). However, evidence for impaired apoptosis in thymocytes remains controversial in NOD mice (Kishimoto and Sprent, 2001; Villunger et al., 2003). Indirect evidence for the lack of peripheral self-tolerance was indicated by the robust proliferation of T-cells derived from NOD mice immunized with random self-peptides without the need for antigen challenge *in vitro* (Ridgway et al., 1996). The spontaneous proliferation of autoreactive T-cells from NOD mice reactive to I-A^g7^ presented by syngeneic antigen-presenting cells *in vitro* was also taken as evidence for impaired self-tolerance (Kanagawa et al., 1998). The autoreactive T lymphocytes that escape to the periphery are thought to be deleted by activation-induced cell death (AICD), which ensues in recently activated T-cells restimulated through the TCR (Green et al., 2003). Compromised AICD was noted inconsistently in CD4^+^ and CD8^+^ T-cells derived from NOD mice upon exposure to the immobilized anti-CD3 antibody *in vitro* (Decallonne et al., 2003; Yang et al., 2004). Regardless of these inconsistencies, these findings favor the hypothesis that overall defective peripheral tolerance could contribute to autoimmune diabetes. Importantly, it has not been demonstrated whether restoration of self-tolerance (AICD) could correlate with the attrition of T1D.

We have demonstrated that the administration of the histone deacetylase (HDAC) inhibitor Trichostatin A (TSA) bestowed irreversible protection against T1D in female NOD mice (Patel et al., 2011; Jayaraman et al., 2013). This was accompanied by reduced inflammation of the pancreatic islets, preservation of insulin-producing beta cells, and altered expression of selected genes. Herein we show that TSA-mediated histone hyperacetylation resulted in equal abrogation of the diabetogenicity of CD4^+^ T-cells cultured under Th1 or Th17 polarizing conditions and selective restoration of AICD in CD4^+^ cells. These data lend unprecedented insights into the mechanisms of autoimmune diabetes that have significant implications for devising intervention strategies to treat patients with T1D.

## RESULTS

### The epigenetic drug comparably regulated the T helper subsets in NOD mice

Female NOD mice (80-100%) procured from the Jackson Laboratory routinely developed diabetes around 20-wk of age when housed in our animal facility (Jayaraman et al., 2010; 2013; 2021; Jayaraman and Jayaraman, 2018; Patel et al., 2011). Weekly treatment with a very low dose of TSA between 16 and 24-wks of age bestowed irreversible protection against diabetes (Figure S1), as reported earlier (Jayaraman et al., 2013; 2021; Patel et al., 2011). We also showed that treatment with the vehicle DMSO neither influenced diabetes incidence nor gene expression. Therefore, we compared the untreated NOD mice with those treated with TSA as in our previous studies. We have also shown that adoptive transfer of unfractionated whole splenocytes from acutely diabetic mice induced diabetes in 100% of the immunodeficient NOD.*scid* mice, which was abolished by TSA treatment (Jayaraman et al., 2013). Consistently, transfer of 2 x 10^7^ T-cell mitogen-activated splenocytes, which contained >98% CD3^+^ cells, as assessed by flow cytometry, transferred diabetes into NOD.*scid* mice (Figure 1). Histone modifier treatment substantially diminished diabetes transferring ability of T-cells to immunodeficient mice.

**Figure 1.**
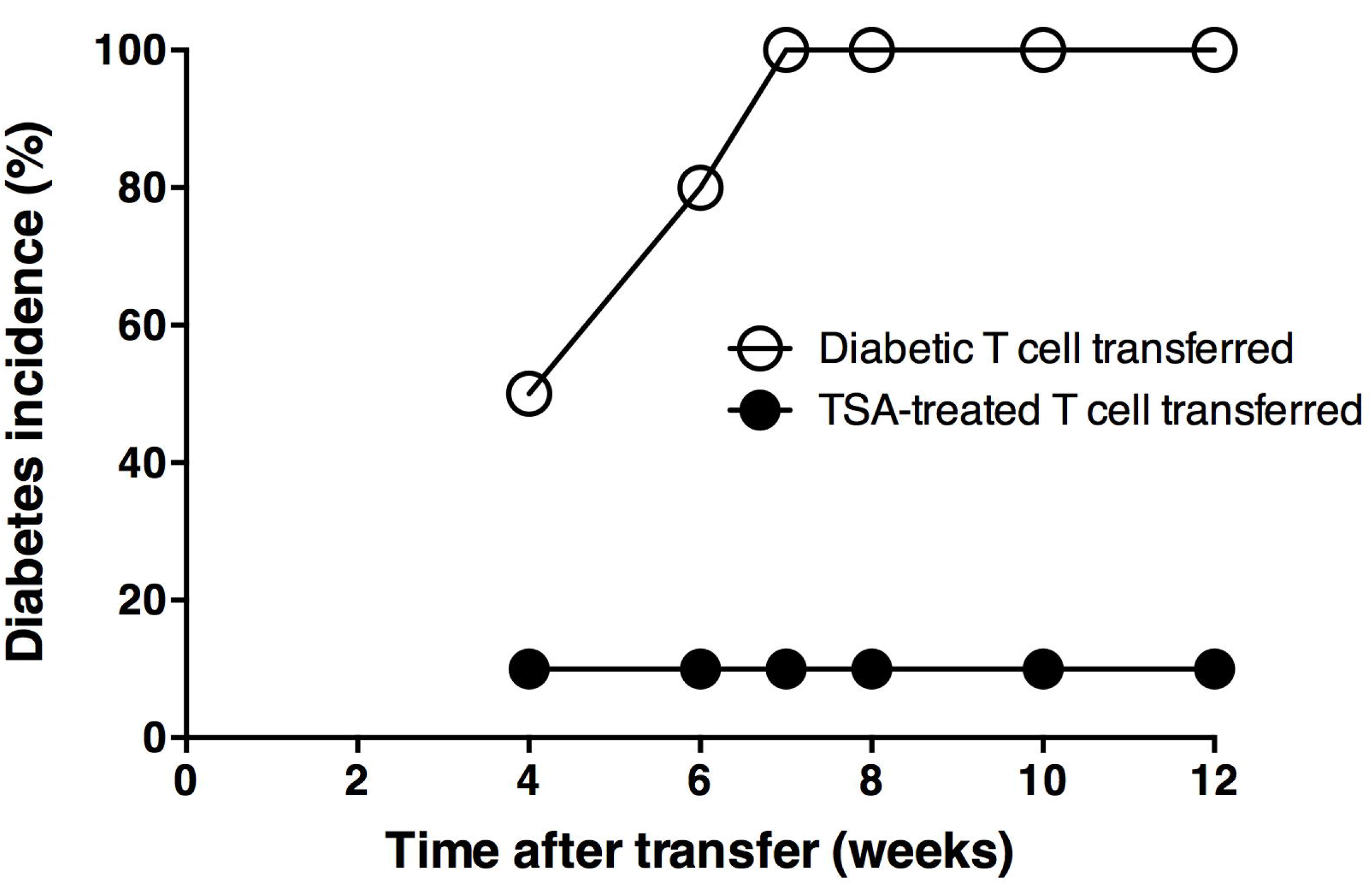
TSA treatment diminished diabetes transferring ability of T lymphocytes. Spleen cells of 28-36 wks old diabetic mice and similarly aged mice treated with TSA from 18 to 24-wks of age were activated Concanavalin A for two days. More than 95% of cells were CD3^+^ as assessed by flow cytometry. The T-cells (2 x 10^7^) from diabetic (empty circles) and TSA-treated mice (filled circles) were injected i.v into NOD.*scid* mice and diabetes monitored. n=10 mice per group.

Previously we have shown that TSA treatment of prediabetic female NOD mice modulated the global gene expression profile of the uninduced whole splenocytes as assessed using an unbiased microarray approach (Jayaraman et al., 2013). Herein, we focused on the effects of histone hyperacetylation on CD4^+^ T lymphocytes that cause diabetes in this model. To this end, we analyzed the frequency of lymphokine-expressing CD3^+^ cells in the spleens of 28-36-wks old diabetic mice and their modulation by TSA treatment. Total splenocytes were stimulated with pharmacological reagents that mimic TCR-signaling, PMA + ionomycin and brefeldin A was added to trap lymphokines in the Golgi apparatus. Cells were stained with fluorescently labeled anti-CD3 antibody, permeabilized, and stained with fluorescent antibodies against lymphokines. The CD3^+^ cells were gated and analyzed for the expression of lymphokines intracellularly. The data shown in Figure 2a indicate that almost equal numbers of T-cells producing IL-17A and IFN-γ were evident in the spleens of diabetic mice. Treatment with TSA comparably reduced these numbers. A small fraction of T-cells from diabetic mice produced IL-10, characteristic of T regulatory cells (Gagliani et al., 2015), which were not influenced by TSA treatment.

**Figure 2.**
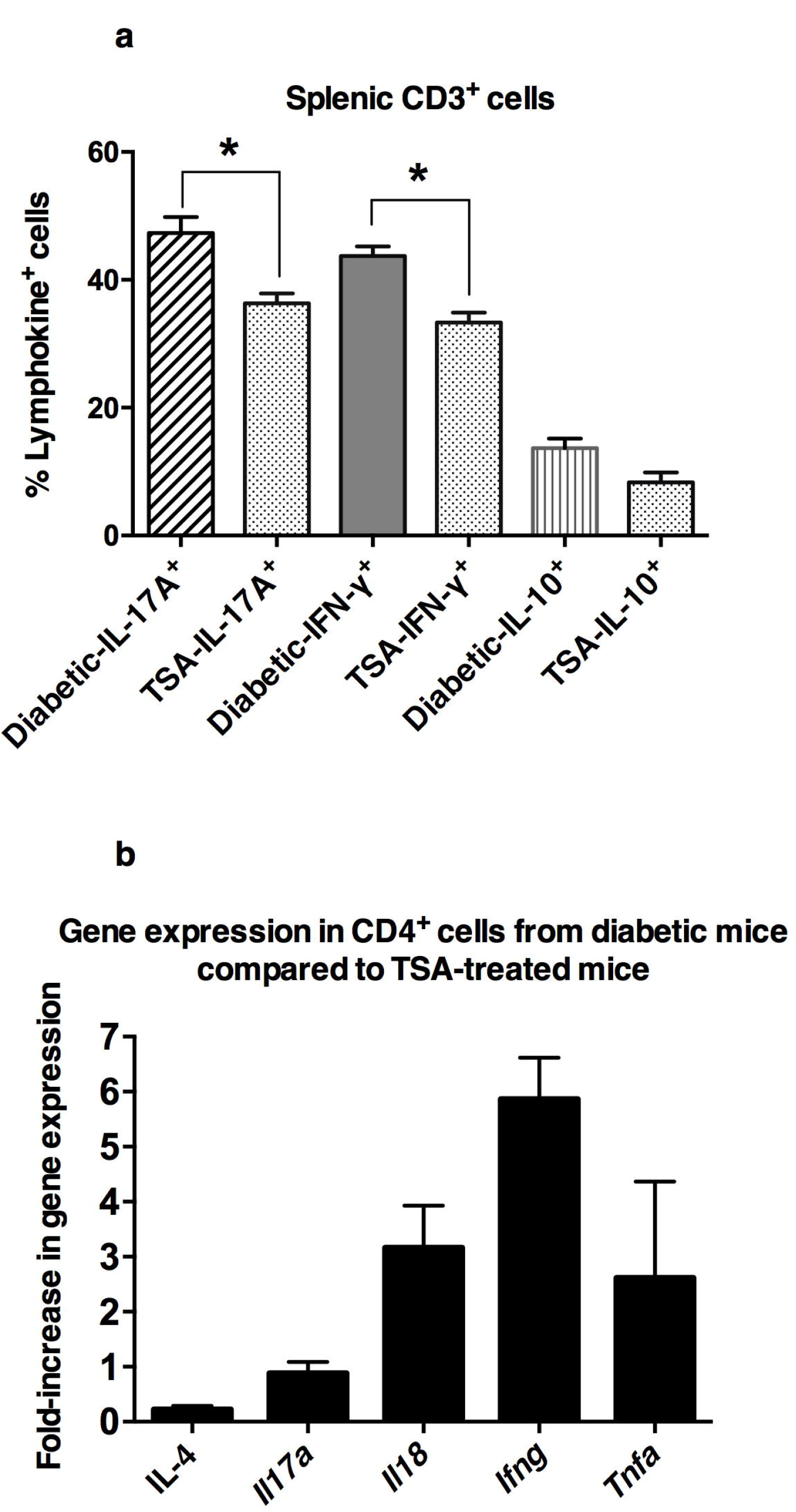
Epigenetic drug regulated the T helper subsets in NOD mice. **a.** Spleen cells from 28-36-wks old diabetic mice and similarly aged mice treated with TSA from 18 to 24-wks of age were activated with PMA and ionomycin in the presence of brefeldin A. Intracellular expression of IFN-γ, IL-17A, and IL-10 was analyzed by staining with anti-CD3 antibody followed by permeabilization and incubation with indicated antibodies conjugated with various fluorochromes. CD3-gated cells were analyzed for lymphokine expression by flow cytometry. Mean +/- SEM of 5 mice per group are shown. Asterisks indicate significant differences (*P*<0.05). **b.** Total RNA was extracted from *unstimulated* CD4^+^ T-cells derived from diabetic and TSA-treated mice, as indicated above. *Basal level* expression of indicated genes was assessed by qRT-PCR in comparison to *Gapdh*. The difference in fold-increase in expression between diabetic and drug-treated mice is shown. Five mice per group were examined in triplicate. Representative data from two different experiments are shown.

The CD4^+^ T lymphocytes isolated from 28-36 wks old diabetic mice and those treated with TSA were stimulated *in vitro* with immobilized anti-CD3 antibodies to induce gene transcription. Analysis of CD4^+^ cells from diabetic mice by qRT-PCR revealed higher expression of *Ifng*, *Il18,* and *Tnfa* while the level of *Il17a* was modest compared to drug-treated cells (Figure 2b). These data are consistent with the possibility that downregulation of both IFN-γ and IL-17A is associated with the abrogation of diabetogenicity of *wild-type* CD4^+^ T cells.

### Polarized *wild-type* Th1 and Th17 cells equally transferred diabetes to NOD.*scid* mice

The Th1 cells polarized *ex vivo* using the BDC2.5 transgenic T cells could adoptively transfer diabetes into neonatal NOD mice (Katz et al., 1995), but polarized Th17 cells failed to do so (Martin-Orozco et al., 2009). Interestingly, the transferred Th17 cells caused diabetes in immunodeficient NOD.*scid* mice, presumably only after conversion into Th1 cells (Martin-Orozco et al., 2009, Bending et al., 2009). It is essential to verify whether this is unique to the transgenic T cells or the Th17 cells from the *wild-type* NOD mice also are required to transdifferentiate into Th1 cells to cause diabetes. To this end, CD4^+^ T cells were isolated from the spleen and lymph nodes of 12-wk old, *prediabetic mice* (as indicated by normal blood glucose level) and cultured under standard Th1 and Th17 polarizing conditions described earlier (Martin-Orozco et al., 2009, Bending et al., 2009). At the end of 8-10 days of culture, cells were washed and re-cultured in anti-CD3 coated plates overnight. Assessment of lymphokines using ELISA indicated that IL-17A was exclusively produced when T cells cultured under Th17 polarizing conditions were stimulated through the TCR (Figure 3a). As expected, similarly activated Th1 cells produced copious amounts of IFN-γ. Unexpectedly, both undifferentiated Th0 cells and those cultured under Th17 polarizing conditions expressed low but detectable levels of IFN-γ. All three types of T helper cells expressed negligible amounts of IL-4 and TNF-α. Analysis by qRT-PCR confirmed the predominant transcription of *Ifng* and modest expression of *Il4, Il17a, Il18,* and *Tnfa* in Th1 cells polarized from the *wild-type* mice compared to Th17 cells (Figure 3b).

**Figure 3.**
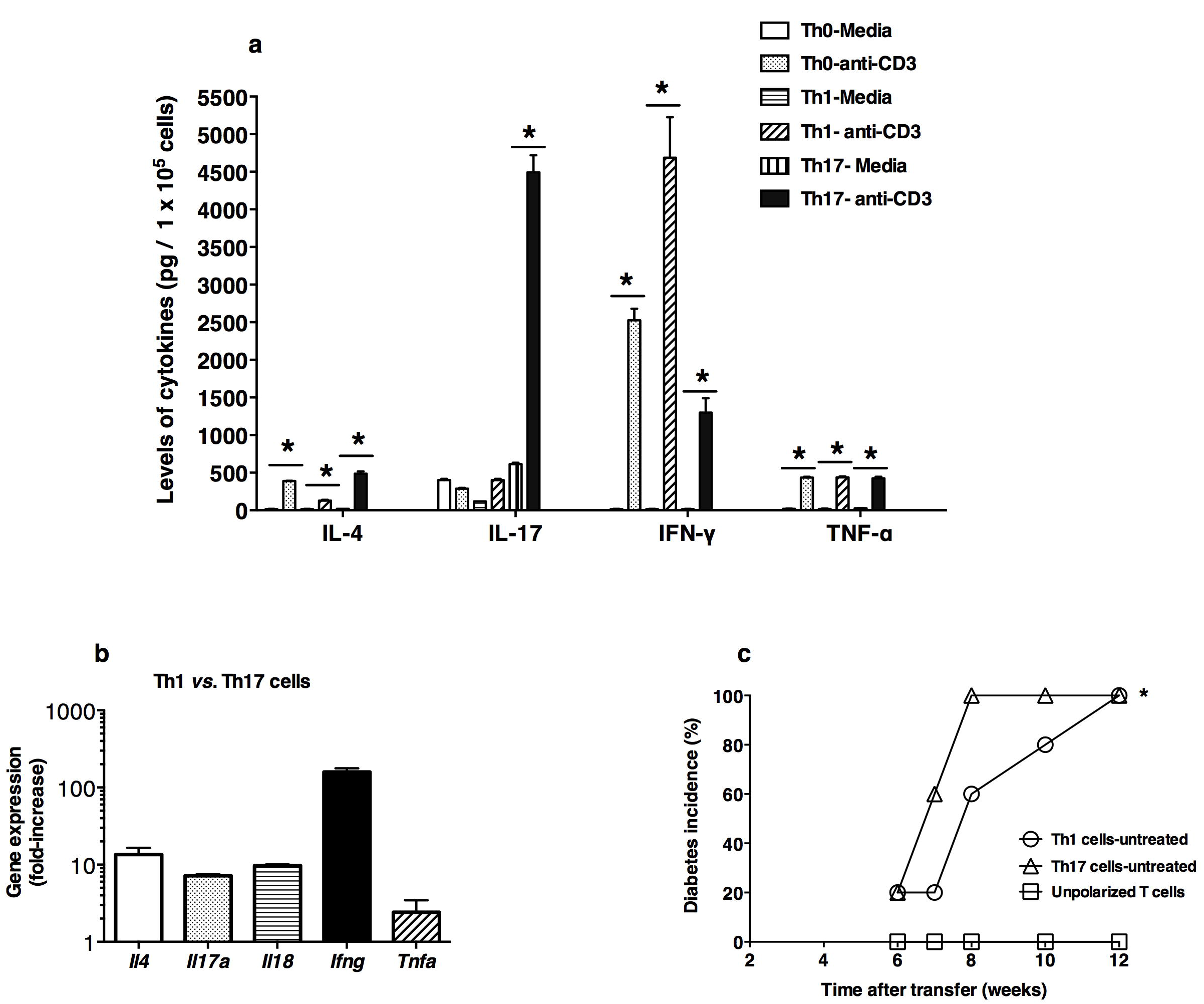
The Th1 and Th17 cells polarized from wild-type NOD T cells transferred diabetes into NOD.*scid* mice. **a.** Purified CD4^+^ T cells derived from *prediabetic* (12-16-wks old) mice were cultured under Th0, Th1, and Th17 polarizing conditions for 8-10 days. At the end of the culture period, cells collected, washed, and incubated on plates precoated with the anti- CD3 antibody. The supernatant was collected after 18 hr and assayed for the presence of lymphokines by ELISA. Data shown are mean +/- SD of triplicate determinations. Representative data from two different experiments are shown. Statistical significance (*P*<0.05) between cells incubated in media and those exposed to the immobilized anti- CD3 antibody is shown by an asterisk. **b.** Gene expression was analyzed in Th1 and Th17 cells polarized from *prediabetic* (12-16-wks old) mice by qRT-PCR in triplicate. The fold-increase in gene expression is depicted. Data are representative of three different experiments. **c.** CD4^+^ T-cells were purified from *prediabetic* (12-16-wks old) mice and cultured under Th0 (empty rectangle), Th1 (empty circle), and Th17 (empty triangle) polarizing conditions. Cells (1. 5 x 10^6^ cells) were injected into NOD.*scid* mice and diabetes was monitored weekly. Diabetes was defined as having >250 mg/dL of glucose in two consecutive weeks. Data were pooled from two experiments consisting of 5 mice per group.

We ascertained the diabetogenic potential of polarized *wild-type* T helper subsets by transferring 1.5 x 10^6^ cells i.v into immunodeficient NOD.*scid* mice. Adoptive transfer of Th1 or Th17 cells induced diabetes in 100% of immunodeficient recipients by 12-wk of cell transfer (Figure 3c). The unpolarized Th0 cells cultured with immobilized anti- CD3 antibody and IL-2 without the differentiating lymphokines failed to transfer diabetes. These data indicate that T-cells require either the Th1 or Th17 polarizing conditions in addition to TCR-mediated activation to express the diabetogenic potential.

### Transfer of *wild-type* Th1 or Th17 cells into NOD.*scid* mice resulted in equal amounts of IFN-γ- and IL-17A-expressing cells

Previous studies showed that the Th17 cells polarized from the BDC5.2 TCR transgenic mice transferred diabetes into NOD.*scid* mice, and this was accompanied by the presence of a small number of Th1 cells in the spleen of recipients (Martin-Orozco et al., 2009, Bending et al., 2009). Therefore, we next examined whether the transfer of polarized Th17 cells from *wild-type* NOD mice could convert into IFN-γ-producing Th1 cells to cause diabetes in NOD.*scid* recipients. To this end, 1.5 x 10^6^ Th1 or Th17 cells polarized from prediabetic CD4^+^ cells were transferred i.v into immunodeficient NOD.*scid* mice. The spleens of recipient mice were harvested 13-wks after the cell transfer and analyzed for intracellular cytokines at the single-cell level by flow cytometry following activation with PMA + ionomycin in the presence of brefeldin A. As expected, a proportion of splenocytes from Th1 cell transferred NOD.*scid* mice displayed IFN-γ in response to stimulation (Figure 4a). A small proportion of cells expressed IL-17A alone or with IFN-γ (double-producers). Spleens of immunodeficient mice receiving wild-type Th17 cells contained cells expressing IL-17A, IFN-γ and double-producers (Figure 4b). Interestingly, TNF-α-expressing cells were abundant in NOD.*scid* mice transferred either with Th1 or Th17 cells. Analysis of the pancreas by qRT-PCR indicated abundant transcription of IFN-γ in the recipients of Th1 cells (Figure S2). Interestingly, the *Il17a* transcript was prominently detected in the pancreas of Th17 cell recipients as well. Collectively, these data indicate that the transfer of either Th1 or Th17 cells polarized from the *wild-type* NOD mice resulted in the comparable generation of IFN-γ^+^ and IL- 17A^+^ cells in a lymphopenic environment.

**Figure 4.**
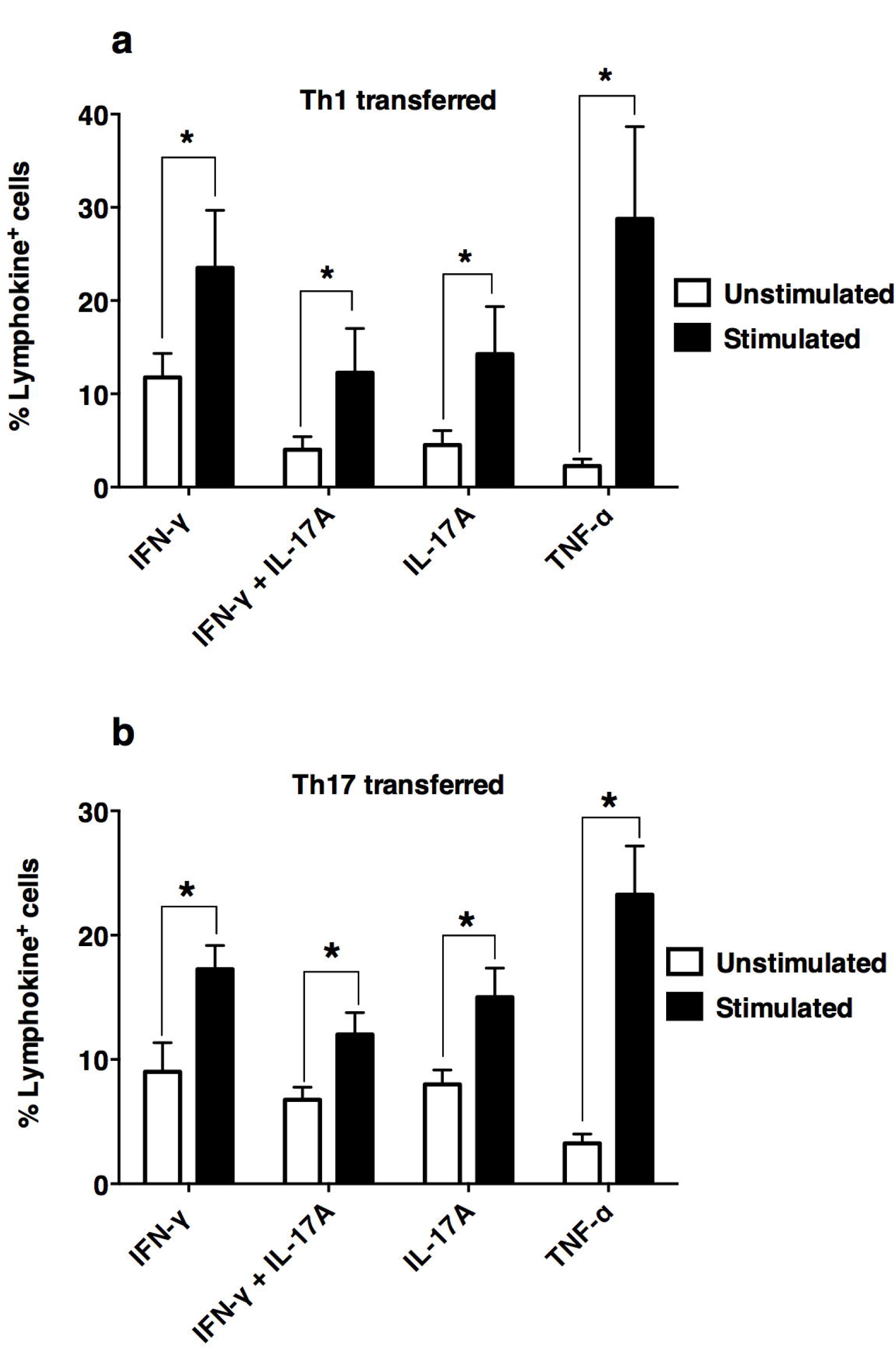
Wild-type Th1 and Th17 cells remained pliable in a lymphopenic environment. NOD.*scid* mice were sacrificed after 13-wks of Th1 (**a**) and Th17 cell (**b**) transfer, as shown in Figure 3c. Splenocytes were assessed for intracellular expression of cytokines by flow cytometry, as described in Figure 2a. Spleens were pooled from five mice per group of NOD.*scid* mice transferred with Th1 or Th17 cells and activated with PMA and ionomycin in the presence of brefeldin A and assayed in triplicate. Data are pooled from two different experiments. Statistical differences (*P*<0.05) between unstimulated and stimulated samples are indicated.

### Histone modifier diminished the diabetogenicity of *wild-type* Th1 and Th17 cells accompanied by repression of signature genes

We next investigated whether altered expression of signature lymphokines in T cells cultured under distinct polarizing conditions accompanied the abrogation of diabetogenicity by the HDAC inhibitor treatment. Transfer of 1. 5 x 10^6^ polarized Th1 cells from diabetic mice induced the disease in 100% of the recipient mice by 12-wks (Figure 5a), consistent with the data shown in Figure 3c. Notably, the same numbers of Th1 cells polarized from TSA-treated mice could transfer diabetes only in 25% of NOD.*scid* mice, indicating a marked reduction in diabetogenicity by drug treatment. Mice were killed after 12-wks of the cell transfer, and the splenocytes of recipients were analyzed for gene expression by qRT-PCR. Consistent with the data presented above (Figure 3a-b), splenocytes from recipients of diabetic Th1 cells upregulated *Ifng* transcription when stimulated with immobilized anti-CD3 antibody *in vitro* (Figure 5b). Interestingly, TSA treatment of T cell donors blunted the ability of Th1 cells to express *Ifng*. As expected, *Il17a* expression was not upregulated in anti-CD3 antibody stimulated NOD.*scid* spleen cells receiving the polarized Th1 cells. These data indicate that the polarized *wild-type* Th1 cells could induce diabetes in a lymphopenic environment without transdifferentiation into Th17 cells. Notably, epigenetic regulation abrogated the diabetogenicity of T-cells cultured under Th1-polarizing conditions.

**Figure 5.**
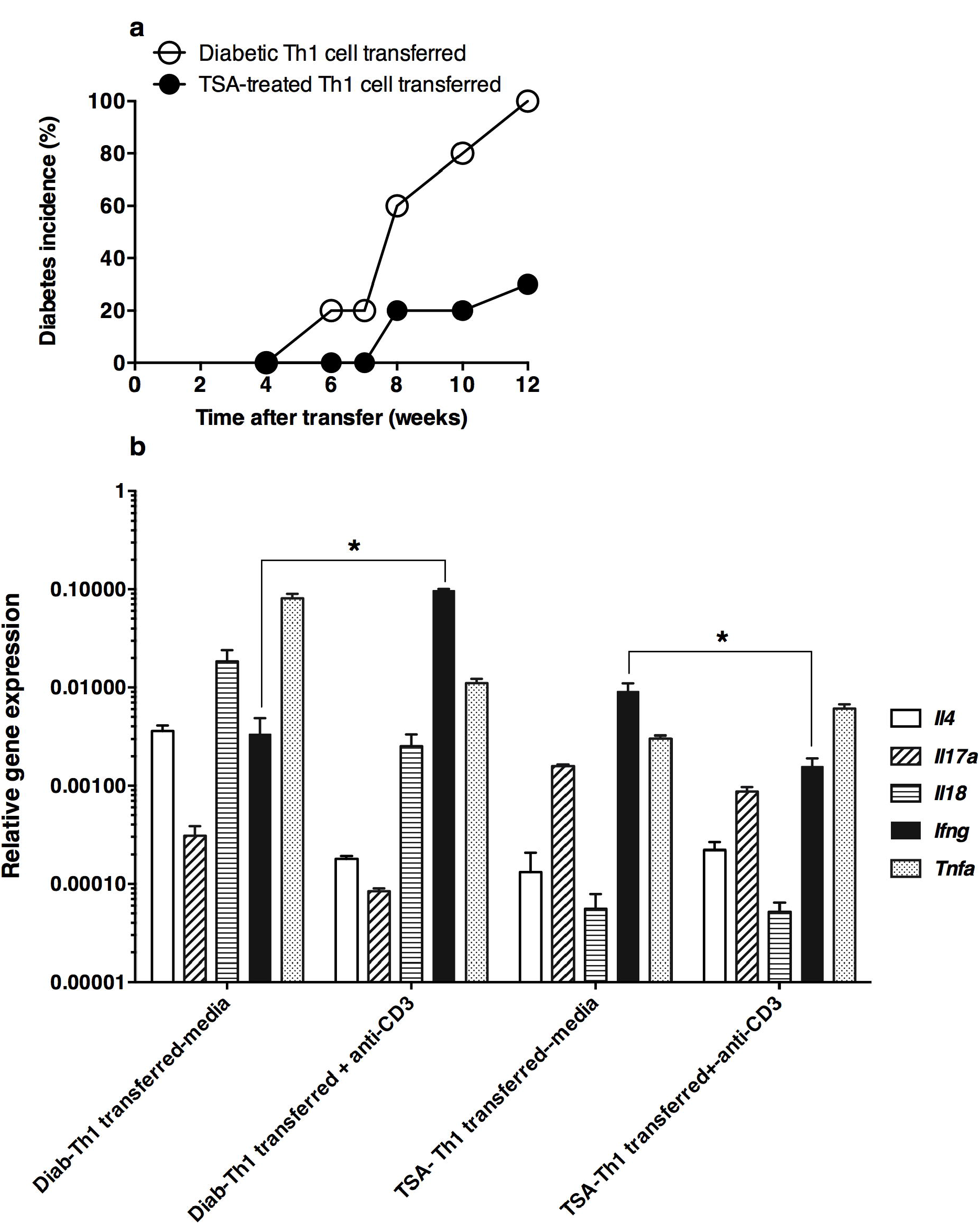
Histone modifier diminished diabetogenicity of wild-type Th1 cells accompanied by altered gene expression. **a.** Splenic CD4^+^ T-cells were purified from five 28-36-wks old diabetic mice (empty circles), and similarly, aged mice treated with TSA (n=5) from 18 to 24-wks of age (filled circles) and cultured under Th1 polarizing conditions. Cells (1. 5 x 10^6^ cells) were transferred to NOD.*scid* mice (5 mice per group), and diabetes monitored weekly. Representative data from two independent experiments are shown. **b.** NOD.*scid* mice were sacrificed after 13 weeks of cell transfer as indicated above. Spleen cells from individual mice were cultured for 18 hr in media alone or in plates precoated with the anti-CD3 antibody. Total RNA was extracted from individual cultures and analyzed for the expression of indicated genes by qRT-PCR in triplicate. Asterisks indicate a statistical difference (*P*<0.05) between unstimulated and stimulated cells. Similar data were obtained in another independent experiment.

Like the Th1 cells, Th17 cells polarized from prediabetic mice transferred diabetes to 100% of the NOD.*scid* mice by 11-wks (Figure 6a). However, Th17 cells polarized from drug-treated, non-diabetic mice transferred diabetes only to 20% of the recipients. The *steady-state level* expression of *Il4*, *Ifng, Il17a*, and its transcription factor *Rorgt* remained higher in mice receiving diabetic Th17 cells than the unmanipulated NOD.*scid* mice (Figure 6b). The *basal* levels of *Il17a* and its transcription factor *Rorgt* were lower in mice receiving Th17 cells derived from drug-treated mice, indicating the repressive effect of TSA on the transcription of these genes. However, the transcription of *Il4*, *Ifng,* and *Gata3* did not differ considerably in mice receiving drug-treated cells. In data not shown, we observed that the splenocytes of NOD.*scid* recipients of TSA-treated cells cultured under Th17 polarizing condition expressed a lower level of *Il17a* but not *Ifng* when activated with anti-CD3 antibody. Thus, treatment of the T cell donors with the histone modifier resulted in comparable attrition of IFN-γ and IL-17A transcription and diabetogenicity.

**Figure 6.**
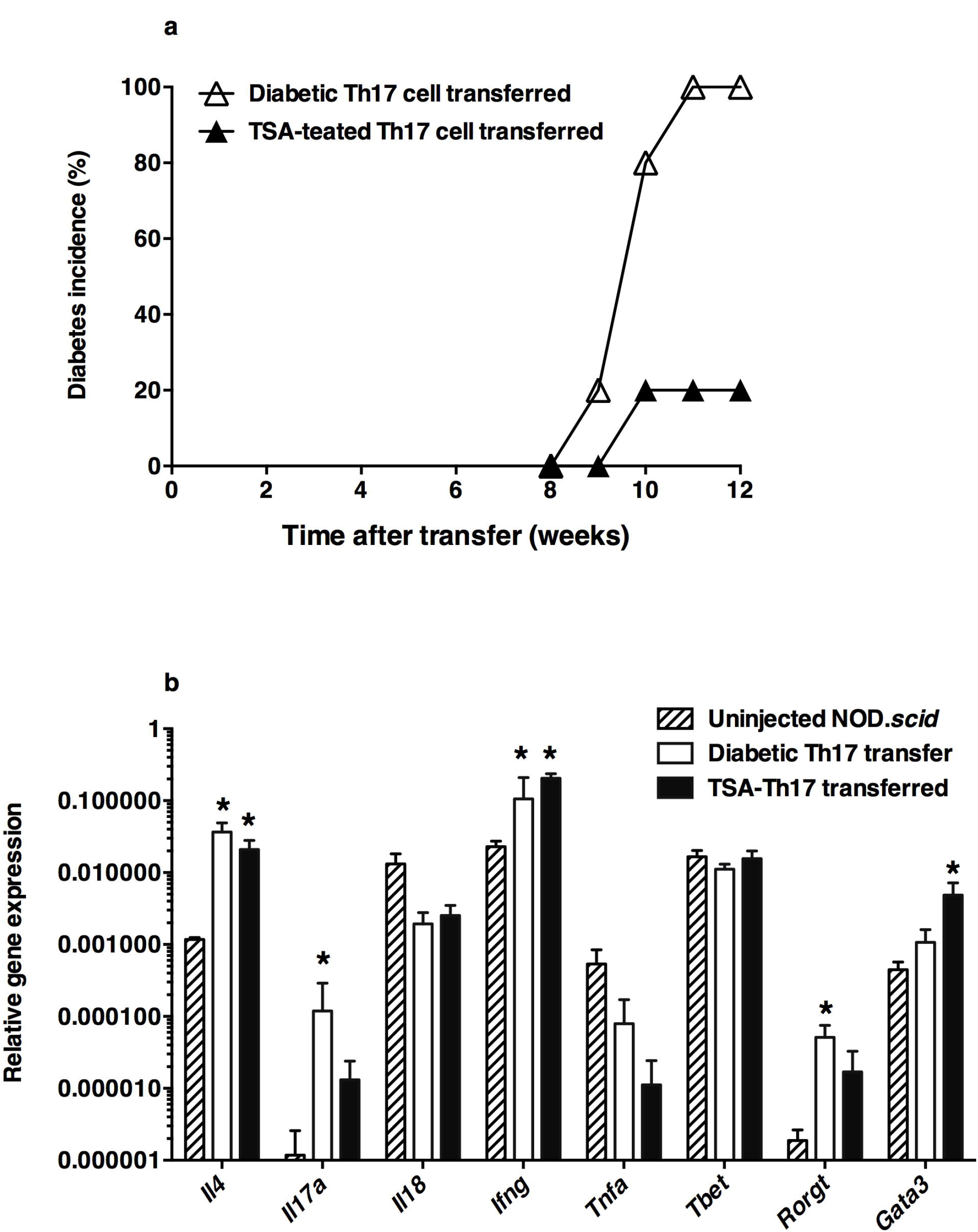
Histone modifier diminished diabetogenicity of wild-type Th17 cells accompanied by altered gene expression. **a.** Polarized Th17 cells obtained from five 28-36-wks old diabetic mice (empty triangles) and similarly aged mice (n=5) treated with TSA from 18 to 24-wks of age (filled triangles). Cells (1. 5 x 10^6^ cells) were transferred to NOD.*scid* mice (5 mice per group) and diabetes monitored weekly. Representative data from two independent experiments are shown. **b.** Gene expression in the spleens of untreated NOD.*scid* mice (hatched bar), and those transferred with Th17 cells derived from diabetic mice (empty bar) or Th17 cells from TSA-treated mice (black bar) by qRT-PCR. Asterisks indicate statistical significance (*P*<0.05) between uninjected mice and those transferred with diabetic or TSA-treated Th17 cells. Five mice per group were investigated. Representative data from two independent experiments are presented.

### TSA treatment restored AICD in CD4^+^ T cells

The AICD mediated by the ligation of the TCR of previously activated T-cells is a potent peripheral tolerance mechanism (Kramer, 1999; Holtzman et al., 2000; Green et al., 2003), and its compromise has been reported in T1D patients (De Maria et al., 1994; Jayaraman et al., 2012). Importantly, it is not clear whether maneuvers that reverse T1D can indeed restore AICD preferentially in the CD4^+^ subset. This critical question was addressed using spleen cells derived from 28-36-wks old diabetic mice and those treated with TSA between 16 to 24-wks of age and rendered diabetes-free. Splenocytes were activated for two days with the T-cell mitogen Concanavalin A, and rHuIL-2 was added to cultures on day three and further cultured for another 3-5 days. At the end of the culture period, >98% of cells were CD3^+^ as assessed by flow cytometry. Enriched T-cells were incubated in plates previously coated with anti-CD3 antibody, and CD4^+^ and CD8^+^ T-cells were enumerated by staining respectively with CD4-FTIC and CD8-PE conjugated antibodies. Dead cells were determined by excluding mBcl, an indicator of intracellular thiols in viable cells (Jayaraman and Jayaraman, 2011). Data shown in Figure 7a indicate that CD4^+^ and CD8^+^ T cells derived from diabetic mice failed to undergo significant apoptosis when activated T-cells were re-exposed to the immobilized anti-CD3 antibody *in vitro*. Significantly, TSA treatment of the T-cell donors restored AICD in CD4^+^ but not CD8^+^ T cells, as indicated by increased cell death following exposure to the immobilized anti-CD3 antibody *in vitro*.

**Figure 7.**
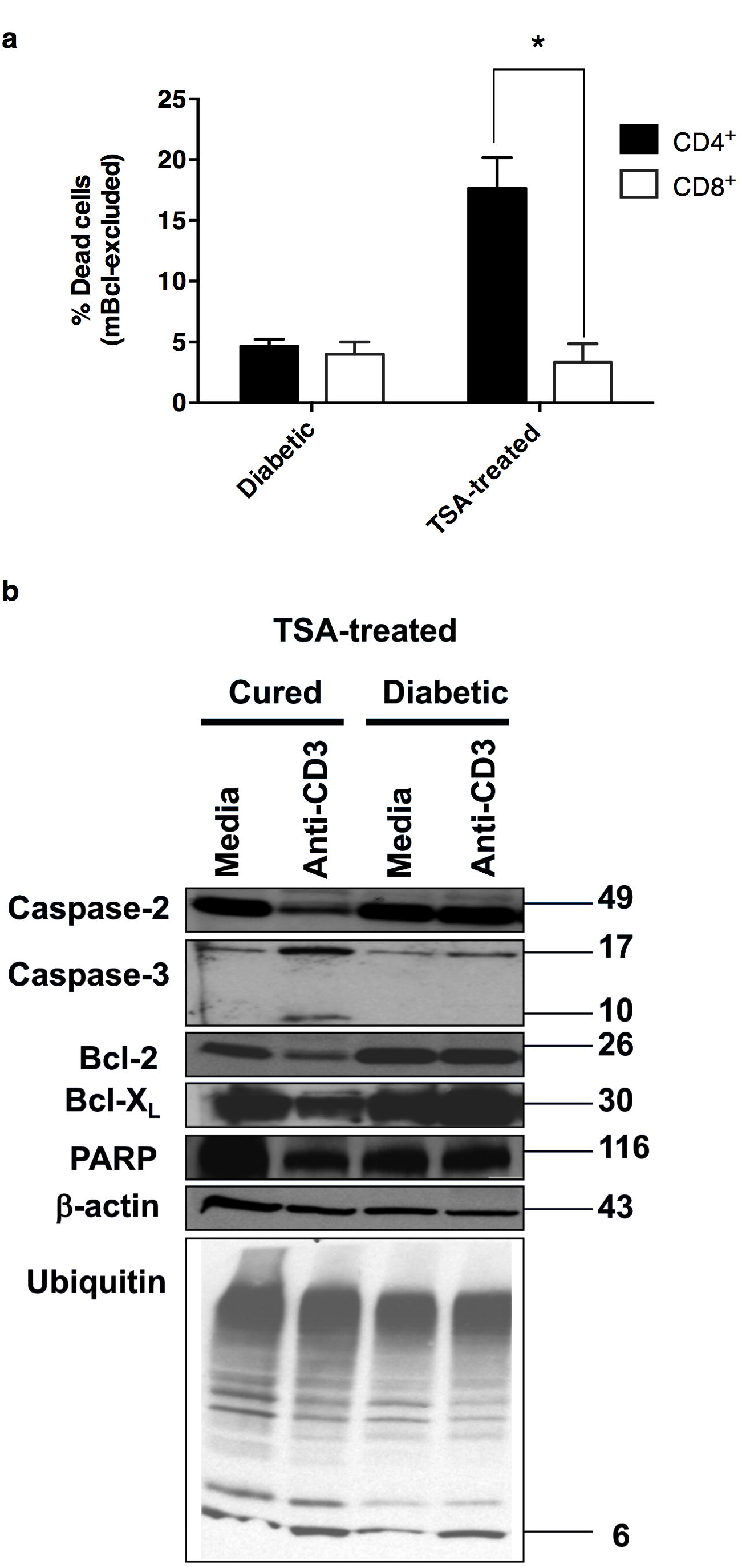
TSA treatment restored AICD in CD4^+^ T cells. **a.** Spleen cells from five 28-36-wks old diabetic mice and similarly aged five mice treated with TSA from 18 to 24-wks of age were cultured individually with Concanavalin A and IL-2 for 3-5 days. Recovered T-cells were incubated overnight on dishes previously coated with the anti-CD3 antibody. Cells were stained with anti-CD4-e450 antibody (filled column) and anti-CD8-FITC (empty column) along with mBcl to monitor glutathione levels as a surrogate marker for viability by flow cytometry. The data are from two independent experiments with similar results. Shown are CD4^+^ and CD8^+^ cells excluding mBcl (dead cells). The asterisk indicates a statistical difference (*P*<0.05) between CD4^+^ T-cells from diabetic and TSA-treated mice. **b.** Spleen cells from diabetic and TSA-treated mice (n=5/group) were cultured with Concanavalin A and IL-2 as described above. Recovered T-cells were cultured overnight on plates previously coated with anti-CD3 antibody or in media alone. Cytosolic proteins were analyzed by western blotting. Indicated are molecular weights of proteins/peptides analyzed, and the difference in the intensity of the bands was confirmed by densitometric analysis using the ImageJ software (not shown). Representative data from two to three independent experiments are shown.

AICD-resistant T lymphocytes from NOD mice displayed lower expression levels of genes implicated in apoptosis execution such as Fas and Fas ligand while increasing the level Bcl-X_L_, a pro-survival factor (Decallonne et al., 2003). To gain insights into the mechanisms of restitution of AICD by the histone modifier, we analyzed the levels of proteins involved in the execution of apoptosis and protection against the extrinsic apoptosis mediated through the TCR (Krammer, 1999; Holtzman et al., 2000; Green et al., 2003). Western blot analysis revealed that the TCR ligation of diabetic T-cells failed to proteolytically activate the upstream caspase-2, the executioner caspase-3, and PARP (Figure 7b). Also, the lack of apoptosis correlated with unaltered levels of pro-survival factors such as Bcl-2 and Bcl-X_L._ In contrast, restoration of AICD by TSA treatment was characterized by the cleavage of caspase-2 and caspase-3 and reduction in anti-apoptotic factors such as Bcl-2 and Bcl-X_L_, and PARP without affecting the ubiquitin level. Densitometric determination using the ImageJ software corroborated the altered levels of these proteins indicated by western blotting (not shown). These data demonstrate that restitution of AICD specifically in CD4^+^ T-cells by TSA treatment engaged the caspase-dependent extrinsic apoptotic pathway.

## DISCUSSION

The AICD is a powerful mechanism that keeps the autoreactive T lymphocytes escaping the thymic deletion under check in the periphery. Evidence for defective self-tolerance in patients with T1D comes from the observation that the peripheral blood T cells displayed aberrant CD3-mediated apoptosis (De Maria et al., 1994; DeFranco et al., 2001; Jayaraman et al., 2012). Although diminished Fas/CD95 expression (Giordano et al., 1995) and Fas-mediated apoptosis (DeFranco et al., 2001) were noted in some T1D patients, we have observed robust apoptosis induced by Fas receptor ligation in long-standing type 1 diabetes patients (Jayaraman et al., 2012). These data are congruent with the notion that the TCR/CD3- and Fas-mediated apoptotic pathways are variously regulated under autoimmune conditions. Similar segregation of these two apoptotic pathways is preserved through the phylogeny. Thus, the expression and apoptotic function of the Fas receptor remained unperturbed in NOD T-cells (Qin et al., 2004) in the face of aberrant TCR/CD3-mediated apoptosis in both the CD4^+^ and CD8^+^ subsets (Figure 7a; Decallonne et al., 2003; Yang et al., 2004). We have observed that TSA treatment of NOD mice bestowed self-tolerance selectively in CD4^+^ cells (Figure 7a), implicating that this could be involved in controlling of T1D as CD4^+^ T-cells are necessary and sufficient to mediate autoimmune diabetes in NOD mice (Christianson et al., 1993). To our knowledge, defective TCR/CD3-mediated apoptosis has not been restituted in T1D patients or NOD mice by maneuvers that reverse autoimmune diabetes. For the first time, we show that TSA treatment rendered NOD mice free of diabetes, which correlated with the restoration of TCR/CD3-mediated apoptosis via the extrinsic pathway. This included proteolytic activation of upstream caspase-2 and -3, degradation of pro-survival factors, Bcl-2 and Bcl-X_L_, as well as the caspase-3-dependent cleavage of PARP (Figure 7b). These findings suggest that the restoration of self-tolerance in CD4^+^ T cells is associated with unmasking or bypassing the upstream blockade of the apoptotic pathway coupled to the TCR/CD3 complex. Further work is necessary to decipher the aberrant signaling elements associated with the TCR/CD3 complex in T lymphocytes of NOD mice (Figure 7a; Decallonne et al., 2003; Yang et al., 2004) and long-standing TID patients (De Maria et al., 1994; DeFranco et al., 2001; Jayaraman et al., 2012).

The sensitivity of functionally distinct T helper subsets such as Th1, Th2, and Th17 to TCR/CD3-mediated apoptosis remains controversial. Whereas the Th17 cells polarized from the hen egg lysozyme-specific TCR transgenic mice were resistant to AICD compared to Th1 cells (Shi et al., 2009), Th17 cells derived from wild-type DBA/1 mice displayed a lower level of AICD than the Th1 and unpolarized Th0 cells (Fang et al., 2010). Although both CD8^+^and CD4^+^ T-cells remained resistant to AICD (Figure 7a), we have not directly determined whether polarized Th1 or Th17 cells could differ in sensitivity to undergo AICD. The CD4^+^ population used in the apoptosis assay consisted of IFN-γ expressing Th1, IL-17A-producing Th17, as well as Th1/Th17 cells which co-expressed IFN-γ and IL-17A (Figure 2a-b; Figure 4a-b). Therefore, it is likely that all of these subsets resist AICD in diabetic mice. Interestingly, TSA treatment restituted self-tolerance in CD4^+^ T-cells (Figure 7b). Concurrently, the histone modifier also abrogated the inducible expression of *Ifng* and *Il17a*, respectively, by Th1 and Th17 cells adoptively transferred into immunodeficient mice (Figure 5b, 6b). These data suggest that sensitivity to AICD and expression of signature lymphokines are inversely correlated. Further work is necessary to decipher the underlying mechanisms linking AICD and turning off T helper cell differentiation *in vitro*.

Although the CD4^+^ T cells (Christianson et al., 1993) and polarized Th1 and Th17 subsets (Bending et al., 2009; Martin-Orozco et al., 2009) adoptively transferred diabetes into NOD.*scid* recipients, little is known about the underlying mechanisms. We have generated Th1 cells from the *wild-type* NOD mice, which expressed IFN-γ but not IL- 17A at the transcriptional and translational levels and readily transferred diabetes into NOD.*scid* recipients (Figure 3a-c). These data support the widely held view that the Th1 cells could mediate diabetes independently of the Th17 cells (Martin-Orozco et al., 2009; Bending et al., 2009). On the other hand, Th17 cells polarized from the transgenic BDC2.5 mice caused pancreatitis and not diabetes in 7-day old NOD mice (Martin-Orozco et al., 2009). Interestingly, the Th17 cells transferred into the immunodeficient mice were thought to mediate diabetes only after conversion to Th1 cells (Martin-Orozco et al., 2009; Bending et al., 2009), implying a subsidiary role for Th17 cells in diabetes. This could be explained by the fact that many Th17 cell preparations included contaminating IFN-γ^+^ cells (Langrish et al., 2005; McGeachy et al., 2007; Martin-Orozco et al., 2009). Consistently, we also detected IFN-γ at the protein and transcript level in polarized *wild-type* Th17 cell preparation (Figure 3a-b). The Th17 cell transfer led to the transcription of both *Ifng* and *Il17a* in the spleen (Figure 6b) and pancreas of NOD.*scid* recipients (Figure S2), indicating that contaminating Th1 cells present in the Th17 preparations could expand in the immunodeficient mice. Nevertheless, the use of similar protocols inadvertently resulted in the existence of a small proportion of Th1 cells in polarized Th17 cells derived from wild-type NOD (this communication), BDC2.5 TCR transgenic (Martin-Orozco et al., 2009), and other strains of mice (Langrish et al., 2005; McGeachy et al., 2007). Thus, the contaminant Th1 cells in Th17 cell preparations expanding in the lymphopenic environment due to homeostatic proliferation and mediating diabetes remains viable.

Although a fraction of Th17 cells also co-expressed IFN-γ (Th1/Th17) in the spleens of NOD.*scid* recipients (Figure 4b), similar to that reported in the EAE models (O’Connor et al., 2008; Jayaraman et al., 2017), their significance in T1D has not been elucidated. These Th1/Th17 cells could relinquish IL-17A expression and differentiate into IFN-γ-producing Th1-like cells in the lymphopenic environment. Since unpolarized Th0 cells also produced IFN-γ (Figure 3a), the contaminating Th0 cells in the Th17 cell preparation may contribute to IFN-γ expression in NOD-*scid* recipients. Some studies suggested reverse plasticity as indicated by the conversion of Th1 to Th17 cells (Liu et al., 2015; Geginat et al., 2016). Although the transfer of Th1 cells resulted in the appearance of IL-17A-producing cells and double-producers in the spleen using the intracellular cytokine assay (Figure 4b), supporting evidence at the mRNA level is lacking (Figure 5b). This anomalous finding suggests that the contaminating Th1/Th17 or Th0 cells in the Th17 preparation could prolong their survival in the lymphopenic environment without actually converting to *Il17a* transcribing cells. In the IL-17 fate mapping mouse strain, Th17 cells partially lost IL-17 expression and upregulated IFN-γ and interestingly, Th1 cells converted to IL-17A/IFN-γ co-expressing cells, indicating that IL-17 expression does not define an end-stage T helper subset (Kurschus et al., 2010). Collectively, these data suggest that various types of T helper cells have the potential to mediate diabetes.

Chromatin remodeling rendered unfractionated splenocytes incapable of transferring diabetes into immunodeficient NOD.*scid* mice (Jayaraman et al., 2013). Consistently, we show herein that purified T lymphocytes (Figure 1a) and polarized Th1 (Figure 5a) and Th17 cells (Figure 6a) derived from TSA-treated prediabetic mice were severely impaired in their ability to cause diabetes in NOD.*scid* mice. The lack of diabetes was associated with diminished frequency of both IFN-γ^+^ and IL-17A^+^ T-cells in the spleen (Figure 2a) and reduced transcription of *Ifng* and *Il17a* in CD4^+^cells (Figure 2b). The latter observation indicates that treatment of T-cell donors with TSA abolished the differentiation of Th1 and Th17 cells *in vitro*, even when they were cultured under appropriate polarizing conditions. This possibility was further validated by the lack of *Ifng* and *Il17a* transcription in anti-CD3-stimulated splenocytes of NOD.*scid* mice respectively receiving CD4^+^ cells derived from drug-treated donors and cultured under Th1 and Th17 polarizing conditions (Figure 5b, 6b). However, *Ifng* transcription was upregulated in the recipients of polarized Th17 cells from untreated mice and those treated with TSA comparably, indicating that the transcriptional regulation of *Ifng* by histone hyperacetylation is variable in Th1 and Th17 cells. Our results suggest that epigenetic regulation of the genome can abolish differentiation of Th1 and Th17 cells and their diabetogenic potential, a novel finding heretofore unreported. Further studies are required to elucidate whether diabetogenicity and T cell differentiation are causally linked.

Despite intense investigation, little is known about the roles of IFN-γ and IL-17A in diabetes induction/manifestation. Treatment with neutralizing anti-IFN- γ antibody reduced diabetes incidence in NOD.*scid* mice transferred with Th17 cells (Martin-Orozco et al., 2009), indicating a crucial role for IFN-γ in Th17-mediated diabetes. On the other hand, the genetic absence of IFN- γ (Hultgren et al., 1996) or the IFN- γ receptor beta chain (Serreze et al., 2000) failed to influence T1D development while a protective role of IFN-γ was also entertained (Trembleau et al., 2003; Zhang et al., 2012). Anti-IL-17A antibody treatment diminished diabetes incidence in NOD mice (Emamaullee et al., 2009) and NOD.*scid* mice transferred with Th17 cells polarized from the BDC2.5 transgenic mice (Jain et al., 2008). However, others found that the neutralizing anti-IFN- γ but not anti-IL-17A antibody ameliorated diabetes induced by transferring Th17 cells polarized from the same TCR transgenic T-cells into NOD.*scid* mice (Martin-Orozco et al., 2009; Bending et al., 2009). Neither IL-17 silencing could protect NOD mice from T1D (Joseph et al., 2012). The use of the IL-17/IFN- γ receptor double-deficient NOD mice suggested that both IL-17 and IFN-γ signaling may synergistically contribute to the development of diabetes in NOD mice (Kuriya et al., 2013). However, we observed that diabetes protection correlated with the transcriptional repression of *Ifng* and *Il17a*, respectively, in drug-treated T-cells cultured under Th1 and Th17 polarizing conditions (Figure 5b, 6b). These cells resemble *undifferentiated* Th0 cells that cannot mediate diabetes but expressed IFN-γ and not IL-17A (Figure 3a). This finding suggests that silencing the signature lymphokines alone is not sufficient for abrogating diabetogenicity. This could be because histone modification can concurrently influence several events crucial for diabetogenesis and T-cell lineage commitment. Our unbiased transcriptome analysis of uninduced splenocytes indicated that TSA treatment selectively repressed a set of inflammatory genes and concurrently upregulated the genes involved in insulin sensitivity, erythropoiesis, hemangioblast generation, and cellular redox control (Jayaraman et al., 2013). Therefore, similar transcriptome analysis of the pathogenic and non-pathogenic (drug-treated) T-cells cultured under Th1 and Th17 polarizing conditions may provide unique perspectives on the genetic basis of diabetogenicity.

In conclusion, the data presented herein show that TSA treatment of *wild-type* NOD mice led to the intervention of the process of commitment to distinct T helper cell lineages and blunted diabetogenic potential. Epigenetic modulation of the genome also restituted self-tolerance selectively in CD4^+^ cells by employing the extrinsic apoptotic pathway. Further research is needed to elucidate whether the transcriptional repression of IFN-γ and IL-17A respectively in Th1 and Th17 cells is linked to abrogation of diabetogenesis. Some autoimmune diseases, such as T1D and EAE are polygenic syndromes (Jayaraman, 2014; 2018; Jayaraman et al., 2020; 2021). Hence, disease manifestation may be multifactorial and concurrent regulation of multiple elements is required for efficient manipulation of autoimmune diabetes (Jayaraman et al., 2021). Epigenetic modifications provide a tool for analyzing events crucial for the manifestation of autoimmune diabetes (Jayaraman, 2014; 2018; Jayaraman et al., 2021). Further work aimed at understanding the epigenetic regulation of diabetogenicity will answer critical questions about autoimmune diabetes.

## AUTHOR CONTRIBUTIONS

VP and AJ conducted experiments, formally analyzed the data, validated the results, and edited the manuscript. SJ conceived, planned, and investigated the project, acquired funding and resources, supervised others, and wrote the original draft, reviewed and edited.

## METHODS

### Mice and treatment

Experiments were conducted using six to eight weeks old female NOD/ShiLtj (H- 2^g7^) mice purchased from The Jackson Laboratory (Bar Harbor, ME) according to the approved protocol of the University of Illinois at Chicago. Mice were treated s.c. with 500 µg/Kg body weight of TSA (Sigma Aldrich, St. Louis, MO) at weekly intervals between 16 and 24 wks of age (Patel et al., 2011; Jayaraman et al., 2013; 2021). More than 250 mg/dL of non-fasting blood glucose on two weekly determinations was considered diabetic.

### Intracellular cytokine assays

Spleen and lymph node cells (5-10 × 10^6^ /ml) of diabetic and TSA-treated mice and NOD.*scid* mice transferred with Th1 or Th17 cells were cultured with PMA (100 ng), ionomycin (one μg) for 4 hr, as described earlier (Jayaraman et al., 2017). Cells were stained with CD3e eFluor450, fixed, permeabilized, blocked with 10% goat serum, and stained with anti-IFN-γ-FITC (clone XMG1.2), anti-IL-17A-PE (clone eBio17B7, eBioscience), or APC-conjugated monoclonal anti-IL-10 antibody (clone JES5-16E3, eBioscience). Cells were analyzed on a BD flow cytometer using FlowJo 6.3.4 software (Treestar), and typical cytograms were obtained, as shown previously (Jayaraman et al., 2017).

### Gene expression analysis

The CD4^+^ T-cells were purified from 28-36-wks old diabetic and TSA-treated mice. The T-cells cultured under Th0, Th1, and Th17 polarizing conditions were collected after 8-10 days. All T-cells were cultured at 1 x 10^6^ cells/ml in plates coated with five µg/ml of anti-CD3 antibody (145-2C11, eBiocience, San Diego, CA) for 18 hr. The spleen and pancreas of NOD*.scid* mice transferred with Th1 or Th17 cells polarized from diabetic or TSA-treated mice were harvested 12-wks after the cell transfer and challenged overnight with 100 ng/ml of PMA (phorbol myristate acetate, Sigma) and one µg of ionomycin (Sigma). The total RNA was extracted and converted to cDNA. Quantitative real-time PCR (qRT-PCR) was performed in *triplicate* on Applied Biosystems ViiA7 Real-time PCR system using one µl of cDNA equivalent to 100 ng of total RNA and 2X SYBR Green master mix. Primers (DNA Technologies, Coralville, IA) were designed and validated previously (Patel et al., 2011; Jayaraman et al., 2013; 2017; 2020; 2021). The gene expression level in each sample was ascertained using *Gapdh* as the normalizer and the 2^-ΔΔ^T method.

### Polarization of Th subsets *in vitro*

The CD4^+^ T cells were purified from the spleen and lymph nodes of prediabetic 12-16-wks old female NOD mice and diabetic mice and those treated with TSA, using the untouched CD4^+^ T cell isolation kit II (Miltenyi Biotec, Germany). More than 98% of purified cells were routinely CD4^+^, as assessed by flow cytometry following staining with anti-CD4-FITC or anti-CD4-PE (eBioscience). Purified CD4^+^ T cells (1 x 10^6^) were cultured in microplates coated with five µg/ml of anti-CD3 antibody and one µg/ml of anti-CD28 antibody (clone 37.51, eBioscience). The Th0 cells received rHuIL-2 without additional cytokines. For the polarization of Th1 cells, ten µg/ml of anti-IL-4 antibody (11B11, eBioscience), 350 ng/ml of mouse IL-12p70 (eBioscience), and two ng/ml of recombinant mouse IL-12 (R & D Systems, Minneapolis, MN) were added. The Th17 cells were polarized in the presence of 30 µg/ml of anti-IFN-γ (clone XMG 1.2) antibody (eBioscience), 10 µg/ml of anti-IL-4 (11B11) (eBioscience), 30 ng/ml of rIL-6 (R & D Systems), 2 ng/ml of TGFβ (eBioscience), and 10 ng/ml of recombinant IL-23 (R & D Systems). All cultures received rHuIL-2 (20 U/ml, eBioscience) on day one and every 3- 4 days thereafter. After 8-10 days of culture, cells were used for various assays.

### ELISA

Cultured Th0, Th1, and Th17 cells derived from prediabetic mice (1 x 10^6^/ml) were incubated overnight on plates previously coated with five µg/ml of anti-CD3 antibody. The supernatant was assayed for IL-4, IL-17A, IFN-γ, and TNF-α using ELISA Ready-SET Go kits eBioscience (San Diego, CA), as described (Patel et al., 2010; Jayaraman et al., 2017).

### Adoptive transfer of diabetes

Spleens from diabetic 28-36 wks old diabetic mice and similarly aged mice treated with TSA from 18 to 24-wks of age were cultured (10 x 10^6^/ml) with five µg of Concanavalin A (Sigma) for two days (Calderon et al., 2006). Polarized Th1 and Th17 cells and Th0 cells were harvested 8-10 days later. Mitogen stimulated T-cells (2 x 10^7^) and polarized (1.5 x 10^6^) T-cells were injected i.v into individual NOD.*scid* mice and monitored for diabetes.

### Activation-induced cell death (AICD)

Spleen and lymph node cells (5 x 10^6^/ml) harvested from diabetic or mice treated with TSA between 28-36-wks of age were activated with 5 µg/ml of Concanavalin A. After overnight culture, 20 U/ml of rHuIL-2 was added to the cultures and harvested after 3-5 days. Cells were then incubated on plates precoated with five µg/ml of anti-CD3 antibody. After overnight culture, cells were stained with anti-CD4-e450 (eBioscience) and anti-CD8-FITC antibody (eBioscience) followed by incubation with 100 µM mBcl (Monochlorobimane, Invitrogen) as described (Jayaraman and Jayaraman, 2011). The cells (10,000 events) were analyzed on a BD flow cytometer. The data were analyzed using the FlowJo 6.3.4 software (Treestar).

### Western blotting

Cytoplasmic proteins were separated on an SDS-PAGE gel, transferred to PVDF membrane, probed with indicated antibodies, developed with HRP conjugated secondary antibodies, and imaged as we described earlier (Jayaraman and Jayaraman, 2011). Antibodies against β-actin, poly ADP ribose polymerase (PARP), ubiquitin, caspase-2, caspase-3, Bcl-2, and Bcl-X_L_ were obtained from Santa Cruz Biotech (Santa Cruz, CA).

### Statistics

The difference in T1D incidence between controls (untreated and DMSO-treated mice) and those treated with TSA was analyzed for statistical significance using the Wilcoxon Signed Rank Test. The numbers of mice investigated are shown in legends to individual figures. Gene expression data were analyzed for statistical significance between indicated groups by Two-way ANOVA or two-tailed unpaired *t*-test as recommended by the GraphPad Prism (6.0) software (San Diego, CA). A *P*-value of <0.05 was considered significant.

## ACKNOWLEDGEMENTS

Mark Holterman is acknowledged for the support of this work. Rajvir Singh is acknowledged for assistance with western blot analysis.

