## Supplementary data for "Epigenetic reprogramming ameliorates type 1 diabetes by decreasing the generation of Th1 and Th17 subsets and restoring self-tolerance in CD4^+^ T cells"

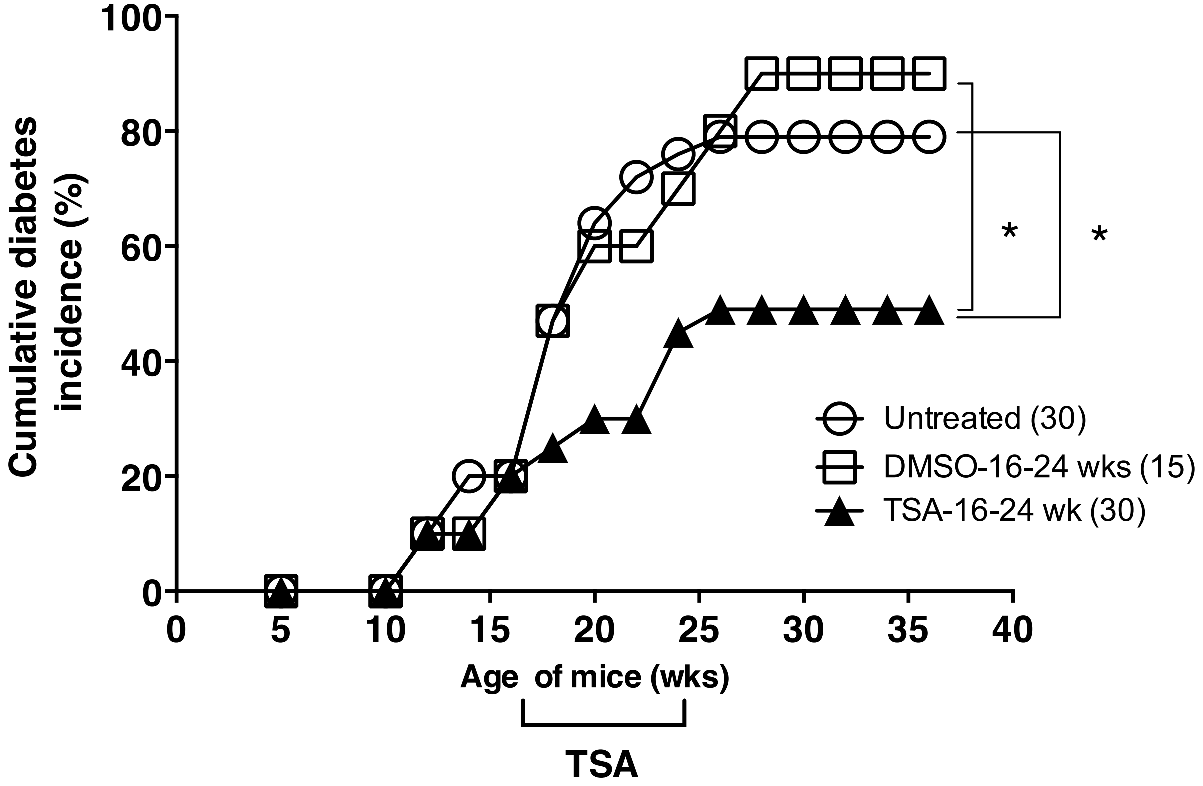


**Supplementary Fig. 1. Treatment with TSA ameliorated diabetes in female NOD mice.** Mice were treated with DMSO or TSA between 16 and 24 wks of age as ndicated. Diabetes was monitored weekly. Numbers of mice investigated are shown in parentheses. Data from three to six experiments were pooled. * *P*<0.03 compared to mice treated with TSA.

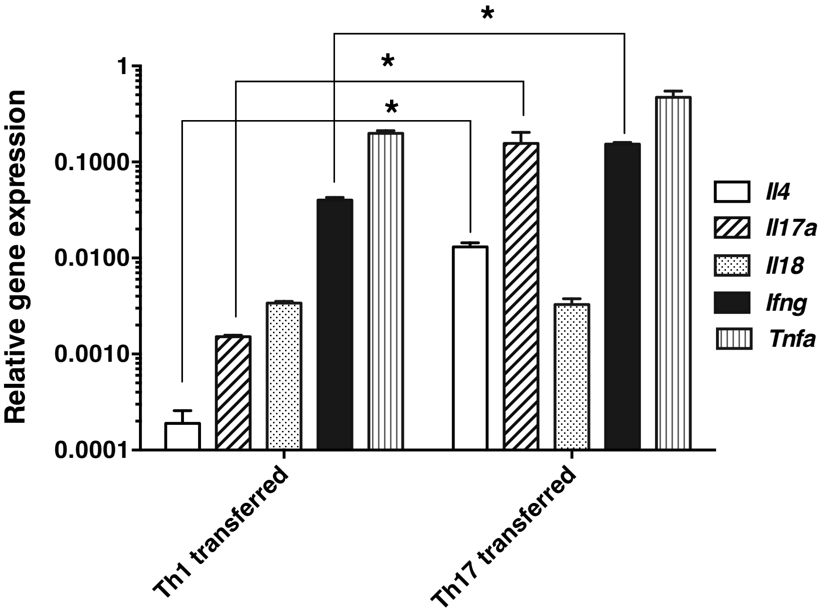


**Supplementary Fig. 2 Gene expression profiles in the pancreas of T helper cell transferred mice.**

Gene expression was analyzed in the pancreas of NOD.*scid* mice receiving Th1 and Th17 cells after 30 days of the cell transfer. n=5-10 mice per group. Statistical significance ($P < 0.05$) is shown between indicated groups by the asterik.
